# Durable Changes to Airway Mucus and Fluid Secretion Following Cholinergic Challenge

**DOI:** 10.1101/2020.03.27.012575

**Authors:** Yan Shin J. Liao, Emily N. Collins, Maria Valentina Guevara, Veronica Schurmann, Kalina R. Atanasova, Laura Bravo, Mariana Sponchiado, Mark J. Hoegger, Leah R. Reznikov

## Abstract

**Purpose:** Mucus abnormalities are central to the pathophysiology of several chronic airway diseases. Mucus secretion and clearance are regulated, in part, by cholinergic innervation. Prolonged cholinergic stimulation may contribute to mucus abnormalities in disease. Thus, we tested the hypothesis that prolonged cholinergic stimulation gives rise to lasting mucus abnormalities in airways.

**Methods:** We delivered aerosolized bethanechol, a cholinergic agonist, to pig airways. Forty-eight hours later, we measured mucus secretion and mucociliary transport in tracheal segments *ex vivo*. Tracheal and bronchoalveolar lavage concentrations of the major secreted mucus glycoproteins, mucin5B (MUC5B) or mucin5AC (MUC5AC), were measured with ELISA and antibody labeling. Pig airway epithelia were cultured at the air-liquid interface and treated with bethanechol for forty-eight hours. Stimulated fluid secretion was measured with reflected microscopy and Ussing chambers were used to measure ion transport.

**Results:** Airways from bethanechol-challenged pigs exhibited sheet-like mucus films, which were not associated with a greater abundance of MUC5AC or MUC5B. Epithelia treated with bethanechol had diminished fluid secretion and decreased Cl^-^ transport. However, mucus and fluid alterations were not associated with impaired mucociliary transport.

**Conclusions:** These data suggest that cholinergic transmission induces sustained alterations in airway mucus properties. Such defects might compound and/or contribute to persistent mucus phenotypes found after the resolution of airway inflammation.

## Introduction

Abnormal mucus composition and clearance highlight several chronic airway diseases, including chronic obstructive pulmonary disease (COPD), cystic fibrosis (CF), and asthma [1–5]. Patients with these diseases often experience periods of stability, punctuated by acute decompensations or exacerbations. These exacerbations are thought to be triggered by noxious environmental stimuli, infection, or allergens, frequently requiring hospitalization. Despite removal from the triggering environment, respiratory compromise is persistent, typically requiring multi-day hospitalization [6, 7]. The mechanism for prolonged airway abnormalities remains unclear, but aberrations in mucus and airway fluid secretion are thought to be critical contributors [8, 9].

Mucus and fluid secretion are regulated by vagal nerve stimulation [10, 11], with airway effector neurons action mediated by the potent secretagogue acetylcholine [11–13]. In turn, the use of anticholinergics as therapeutics in chronic airway disease [14, 15] underscores the importance of the cholinergic system in exacerbation prevention. However, enhanced cholinergic stimulation’s effect on mucus secretion remains unclear, including in the absence of chronic disease. The goal of the current investigation was to determine the sustained impact of enhanced cholinergic stimulation on mucus and fluid secretion. We hypothesize that enhanced cholinergic transmission contributes to abnormal and persistent airway mucus in the absence of airway inflammation.

## Materials and methods

### Animals and ethic statement

A total of 20 male and 20 female piglets (Yorkshire-Landrace, 2-3 days of age) were obtained from a commercial vendor and fed commercial milk replacer (Liqui-Lean). Piglets were allowed a 24-hour acclimation period prior to interventions. Data were collected from 5 separate cohorts of piglets across approximately 1 year. At the conclusion of the experiments, animals were sedated with ketamine (20 mg/kg), and xylazine (2.0 mg/kg), and intravenous propofol (2 mg/kg, Henry Schein Animal Health), followed by euthanasia with intravenous pentobarbital sodium and phenytoin sodium solution (90 mg/kg, Euthasol^®^ Henry Schein Animal Health). The University of Florida Animal Care and Use Committee approved all procedures. Care was in accordance with federal policies and guidelines.

### Airway instillation

After acclimation, piglets were anesthetized with 8% sevoflurane (Sevothesia™, Henry Schein Animal Health). Airways were accessed with a laryngoscope and a laryngotracheal atomizer (MADgic) was passed directly beyond the vocal folds to aerosolize either a 500 μl of 0.9% saline solution (control) or bethanechol chloride in 0.9% saline solution to the airway [16]. The dose selected has previously been shown to acutely increase airway resistance in piglets [16, 17].

### Chemicals and drugs

Bethanechol chloride (Selleckchem) was dissolved in 0.9% saline to final concentration of 8 mg/ml and sterilized with a 0.22 μm filter (Millex GP) [16]. Acetyl-beta-methacholine-chloride (Sigma) was dissolved in 0.9% saline for *ex vivo* application [18]. Drugs used in the Ussing chamber studies: amiloride hydrochloride hydrate (Sigma); carbamylcholine chloride (carbachol; Sigma); forskolin (Sigma); 3-isobutyl-1-methylxanthine (Sigma); Glyh-101 (a gift from the Cystic Fibrosis Foundation Therapeutics and Robert Bridges, Rosalind Franklin University of Medicine and Science, North Chicago, Illinois, USA); and 4,4’-Diisothiocyano-2,2’-stilbenedisulfonic acid (DIDS) disodium salt (Sigma) [19].

### Bronchoalveolar lavage and ELISA

Following euthanasia, the caudal left lung of each piglet was excised and the main bronchus cannulated; three sequential 5 ml lavages of 0.9% sterile saline were administered as previously described [16, 18, 20]. The recovered material was pooled, spun at 500 X g, and supernatant stored at −80°C. A porcine MUC5AC (LSBio, LS-F45847-1) and porcine MUC5B (LSBio, LS-F45852-1) ELISA was performed according to the manufacturer’s instructions in duplicate as previously described [18]. The ELISA was read using a filter-based accuSkan FC micro photometer (Fisher Scientific). The limits of sensitivity were > 0.188 ng/ml and > 0.375 ng/ml for MUC5AC and MUC5B, respectively. The intra-assay and inter-assay coefficient of variability were respectively <6.1% and <5.2% for MUC5AC and <6.5% and <5.9% for MUC5B.

### Immunofluorescence in tracheal cross sections

1-2 tracheal rings were removed post-mortem and embedded in Peel-A-Way embedding molds containing Tissue-Tek OCT (Electron Microscopy Sciences). Molds were placed in a container filled with dry ice until frozen, and subsequently stored at −80 °C long-term. Tissues were sectioned at a thickness of 10μm and mounted onto SuperFrost Plus microscope slides (ThermoFisher Scientific). We used immunofluorescence procedures and analyses similar to those previously described [18, 20, 21]. We used the following anti-mucin antibodies: rabbit anti-MUC5B (1:500; Santa Cruz, Cat. # 20119) and mouse anti-MUC5AC (clone 45M1) (1:1,000, ThermoFisher Scientific, Cat # MA512178).

### Mucus secretion assay *ex vivo*

We measured mucus secretion using previously described methods [18, 22]. Briefly, 3-4 rings of trachea were removed post-mortem and the outside of the tracheas were wrapped in gauze soaked with 5 ml of the following: 135 mM NaCl, 2.4 mM K_2_HPO_4_, 0.6 mM KH_2_PO_4_, 1.2 mM CaCl_2_, 1.2 mM MgCl_2_, 10 mM dextrose, 5 mM HEPES, pH 7.4 (NaOH), 1.5 mg/ml of methacholine, and 100 μM bumetanide [18]. Tracheas were then placed in a 37 °C temperature-controlled humidified incubator for 3 hours. Following stimulation, tracheas were fixed overnight in 4% paraformaldehyde and jacalin-FITC and wheat-germ agglutin-(WGA) rhodamine were used to visual mucus as previously described [18, 22, 23]. Tracheas were imaged with Zeiss Axio Zoom V16 posterior to anterior and scoring assigned as previously described [18].

### *Ex vivo* mucus transport assays

Mucociliary transport was measured using methods similar to those described [18, 24]. Briefly, 3 rings of tracheas were submerged in 5 ml of prewarmed solution containing the following: 138 mM NaCl, 1.5 mM KH_2_PO_4_, 0.9 mM CaCl_2_, 0.5 mM MgCl_2_, 2.67 mM KCl, 8.06 mM Na_2_HPO_4_-7H_2_0, 10 mM HEPES, pH 7.4 (NaOH), and 100 μM bumetanide. Tracheas were placed onto a heated stage and kept at 37 °C. Images were acquired every 1 minute for 35 minutes. After 5 minutes of baseline measurements, methacholine was administered directly into the solution covering the basolateral and apical sides at a dose of 0.004 mg/ml. IMARIS software that utilized principles of the well-validated algorithms published by Jaqaman and colleagues [25] was used to track mucus transport across time measured as previously described [18].

### Primary cultures of differentiated airway epithelia

Epithelial cells were isolated from piglet tracheas by enzymatic digestion (Pronase, Roche; DNase, Sigma) seeded onto collagen (Rat Tail Corning Collagen I, Corning)-coated permeable filter supports (Corning Transwell polycarbonate membrane inserts, area = 0.33 cm^2^, pore size = 4 μm; Corning), and grown at the air-liquid interface using previously described protocols [26]. Differentiated epithelia were studied at a minimum of 14 days after seeding. Medium consisted of DMEM/F12 supplemented with 2% Ultroser G (USG, Crescent Chemical) and antibiotics as previously described [26]. Medium was replaced every 2-3 days. Epithelia were cultured in USG medium containing 10 μM bethanechol/0.9% saline for 48 h. Control epithelia were cultured in USG containing equivalent concentrations of 0.9% saline.

### Electrophysiological measurements of cultured airway epithelia

Cultured epithelia were studied in modified Ussing chambers (EasyMount Ussing Chamber System; Physiologic Instruments) as previously reported [19]. Transepithelial voltage was maintained at 0 mV and short-circuit current (Isc) measured (VCC MC-8; Physiologic Instruments). Isc and delta (Δ) Isc were reported.

Cultured airway epithelia were bathed on the apical and basolateral surfaces with the following solution: 135 mM NaCl, 2.4 mM K_2_HPO_4_, 0.6 mM KH_2_PO_4_, 1.2 mM CaCl_2_, 1.2 mM MgCl_2_, 10 mM dextrose, 5 mM HEPES, pH 7.4 (NaOH). The solution was maintained at 37°C and gassed with compressed air [27, 28].

The following protocol was performed: (a) measurements at baseline; (b) apical 100 μM amiloride to inhibit ENaC; (c) 100 μM basolateral carbachol to simulate calcium-activated chloride channels (d) 100 μM DIDS apical to inhibit calcium-activated chloride channels [29]; (e) 10 μM apical forskolin and 100 μM IBMX to increase cAMP and activate CFTR; and (f) 100 μM GlyH-101 apical to inhibit CFTR. All drug concentrations were selected according to our previous studies [27, 28].

### Fluid secretion studies

We implemented the reflected light microscopy method to measure the meniscus of the secreted fluid as previously described [30]. Briefly, cultures were placed on a heated stage maintained at 37°C and imaged using a Zeiss Axio Zoom V16. The following buffer bathed the basolateral side: 135 mM NaCl, 2.4 mM K_2_HPO_4_, 0.6 mM KH_2_PO_4_, 1.2 mM CaCl_2_, 1.2 mM MgCl_2_, 10 mM dextrose, 5 mM HEPES, pH 7.4 (NaOH). Light intensity and magnification were held constant for all cultures. A single image was taken at designated time points. Cultures were returned to the incubator in between imaging. The intensity profile of the image was examined using Zen Pro software analysis (Zeiss). The XY coordinates of the intensity profile correlated to distance (X) and intensity (Y). The length of the fluid meniscus could therefore be readily identified as the distance between the greatest decrease in pixel intensity (at cell culture insert edge) minus the average baseline intensity (in the middle of the culture). A 4-point regressive curve was used to convert meniscus lengths to fluid volumes. Apical fluid secretion to 100 μM basolateral carbachol was measured.

### Statistical analysis

A two-way ANOVA (sex as one factor, treatment as the other factor) was performed initially to investigate potential sex differences. No significant interactions between sex and treatment were verified for any of the variables studied, indicating that sex did not affect the response to treatments. Thus, separation based upon sex was not strongly justified [18, 31], and therefore we grouped the data to represent the population better. Two-way ANOVA analyses stratified by sex are available in Table 1. A test for sex differences was not statistically valid for fluid secretion assays, as there were only two animals from each sex per group. For parametric data that compared two groups, we used unpaired-test, except for the Ussing chamber studies data, in which cultures were assessed using a paired t-test (+/- bethanechol treatment). For fluid secretion assays, we used a repeated measures ANOVA with Sidak post-hoc comparisons. Non-parametric data were examined by a Mann-Whitney test [16, 18, 20]. All tests were carried out using GraphPad Prism 7.0a. Statistical significance was determined as P < 0.05.

**Table 1.** Results separated by sex. Data are shown as Mean ± SEM.

| Variables | Groups |  |  |  | Two-way ANOVA P-values |  |  |
| --- | --- | --- | --- | --- | --- | --- | --- |
|  | Female |  | Male |  | Sex | Treatment | Sex*<br>Treatment |
|  | Female Saline | Female Bethanechol | Male Saline | Male Bethanechol |  |  |  |
| Bronchoalveolar lavage concentrations of MUC5AC | 0.041 ± 0.005 | 0.035 ± 0.003 | 0.043 ± 0.006 | 0.042 ± 0.005 | 0.3620 | 0.4410 | 0.6285 |
| Bronchoalveolar lavage concentrations of MUC5B | 0.20 ± 0.041 | 0.19 ± 0.035 | 0.23 ± 0.039 | 0.22 ± 0.036 | 0.5006 | 0.8510 | 0.9739 |
| Surface MUC5AC mean signal intensity | 87.58 ± 8.36 | 84.21 ± 12.88 | 82.91 ± 9.12 | 108.89 ± 6.96 | 0.3034 | 0.2461 | 0.1345 |
| Surface MUC5B mean signal intensity | 40.59 ± 7.12 | 35.55 ± 6.88 | 41.61 ± 6.87 | 35.29 ± 6.08 | 0.9554 | 0.4058 | 0.9247 |
| Gland MUC5AC mean signal intensity | 69.47 ± 6.97 | 52.59 ± 5.46 | 64.21 ± 8.36 | 67.99 ± 9.66 | 0.4049 | 0.5183 | 0.1922 |
| Gland MUC5B mean signal intensity | 77.73 ± 10.52 | 72.52 ± 11.39 | 77.97 ± 10.15 | 73.45 ± 10.89 | 0.9564 | 0.6534 | 0.9747 |
| Basal short circuit current | 24.27 ± 3.35 | 27.83 ± 1.13 | 25.90 ± 6.25 | 24.95 ± 6.38 | 0.9115 | 0.8163 | 0.6892 |
| Amiloride-sensitive short current | -8.4 ± 4.11 | -9.07 ± 3.90 | -10.49 ± 2.27 | -7.885 ± 1.30 | 0.8748 | 0.7384 | 0.5755 |
| Carbachol-sensitive short current | 5.09 ± 2.56 | 3.03 ± 1.73 | 2.37 ± 0.41 | 1.4 ± 0.53 | 0.1377 | 0.2888 | 0.6951 |
|  | Female |  | Male |  | Sex | Treatment | Sex*<br>Treatment |
|  | Female Saline | Female Bethanechol | Male Saline | Male Bethanechol |  |  |  |
| DIDS-sensitive short current | -3.89 ± 0.52 | -2.7 ± 0.09 | -5.77± 2.17 | -4.21± 1.65 | 0.3230 | 0.4175 | 0.9117 |
| F&I-sensitive short current | 17.48 ± 5.69 | 15.37 ± 4.96 | 17.56 ± 6.27 | 14.84 ± 6.07 | 0.9717 | 0.6982 | 0.9606 |
| GLYH-sensitive short current | -12.32± 3.70 | -16.37 ±1.93 | -22.85± 9.87 | -22.05± 10.51 | 0.3739 | 0.8557 | 0.7865 |
| Mean jacalin-labeled sheet index scores | 1.16 ± 0.07 | 1.54 ± 0.16 | 1.42 ± 0.14 | 2.15 ± 0.25 | 0.0145 | 0.0023 | 0.3084 |
| Mean WGA-labeled sheet index scores | 1.06 ± 0.04 | 1.22 ± 0.14 | 1.3 ± 0.15 | 1.57 ± 0.23 | 0.0673 | 0.1789 | 0.7368 |
| Mean Jacalin-labeled strand index scores | 1.12 ± 0.1 | 1.14 ± 0.07 | 1.28 ± 0.09 | 1.12 ± 0.06 | 0.3942 | 0.3942 | 0.2749 |
| Mean WGA-labeled strand index scores | 1.02 ± 0.02 | 1.12 ± 0.07 | 1.08 ± 0.03 | 1.08 ± 0.04 | 0.8248 | 0.2723 | 0.2723 |
| Mean speed (µm/s) of mucus transport | 1.28±0.11 | 1.35±0.09 | 1.14±0.11 | 1.18±0.10 | 0.1236 | 0.5672 | 0.8635 |
| Max speed (µm/s) of mucus transport | 5.20±0.46 | 5.36±0.43 | 4.979±0.43 | 5.07±0.46 | 0.5799 | 0.7923 | 0.9385 |
| Mean length (µm) of mucus particle track | 1339.83±181.81 | 1398.57±134.30 | 1210.32±123.02 | 1313.82±140.82 | 0.4699 | 0.5837 | 0.8796 |
|  | Female |  | Male |  | Sex | Treatment | Sex*<br>Treatment |
|  | Female Saline | Female Bethanechol | Male Saline | Male Bethanechol |  |  |  |
| Mean intensity of fluorescently labeled mucus particles on the tracheal surface | 10.98±1.09 | 15.54±2.62 | 16.85±2.58 | 15.00±2.39 | 0.2452 | 0.5535 | 0.1644 |

## Results

We performed multi-level analyses to examine mucus secretion properties. We first investigated mucus morphology and secretion *ex vivo*. We stained mucus with jacalin [22] and WGA[22], surrogates for MUC5AC and MUC5B, respectively [18]. We found that airways from pigs challenged *in vivo* with bethanechol exhibited a statistically significant increase in film-like mucus sheets for jacalin-labeled mucus (Figure 1A-C), but not for mucus labeled with WGA (Figure 1D). Because mucus strands emanating from submucosal glands are characteristic of some airway diseases [22, 32], we also examined tracheal segments stimulated *ex vivo* with methacholine for mucus strand formation [18]. No increases in strand formation for mucus labeled with jacalin (Figure 1E) or WGA (Figure 1F) were found.

**Figure 1.**
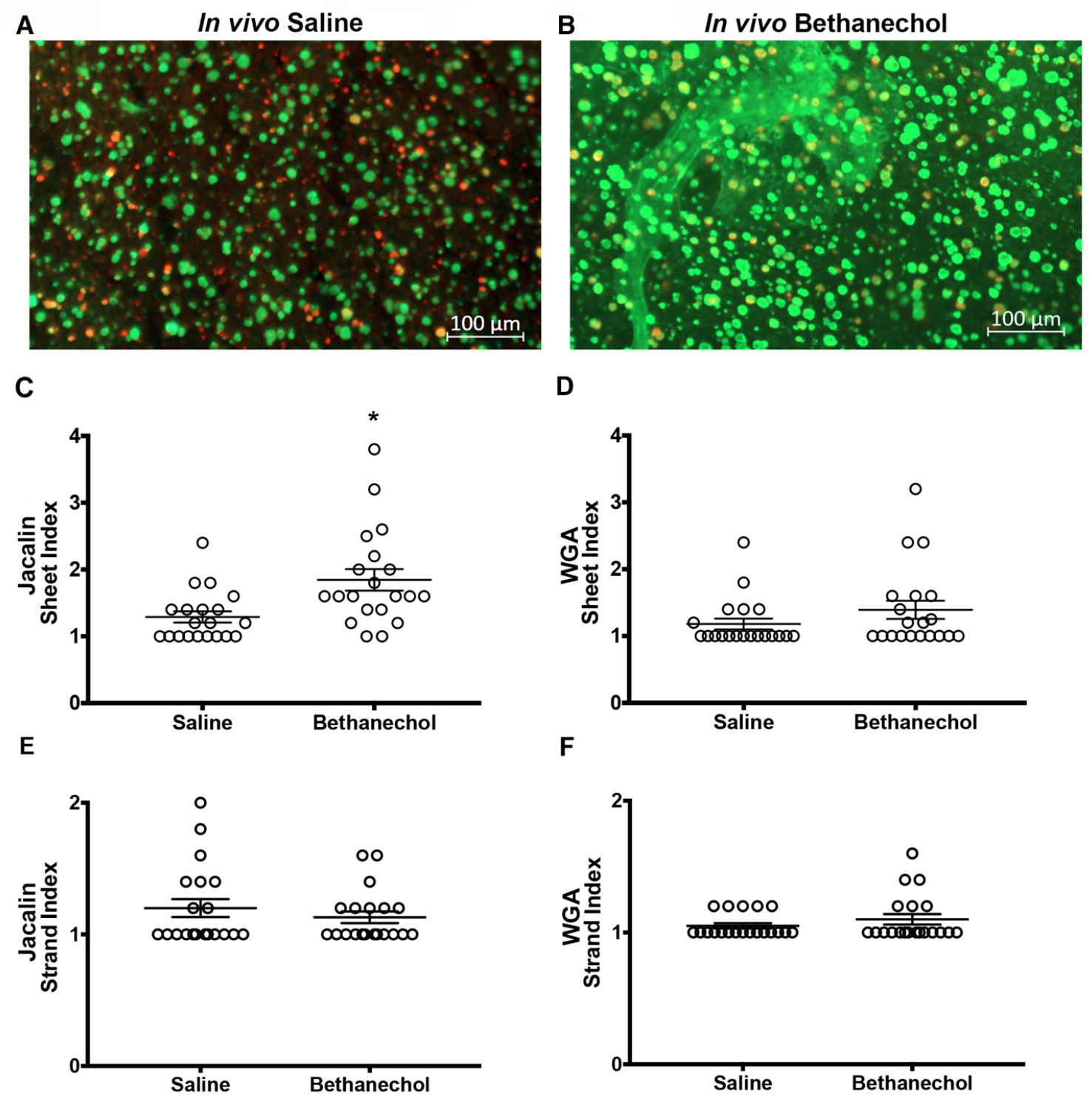
Tracheal mucus morphology in pigs challenged with bethanechol. Representative image of an *ex vivo* trachea from a saline (**A**) or bethanechol (**B**) challenged piglet stimulated with methacholine. Discrete entities of mucus were observed and visualized by jacalin lectin (green) and WGA (red) staining. Sheet index for jacalin-labeled mucus (**C**) and WGA-labeled mucus (**D**). Strand index for jacalin-labeled mucus (**E**) and WGA-labeled (**F**) mucus. n = 20 saline-challenged piglets (10 females, 10 males), n = 20 bethanechol-challenged pigs (10 females, 10 males). Data points represent the mean score for each piglet calculated from 5-7 analyzed images (encompassing the anterior, middle and posterior regions of the trachea). A greater score indicates greater incidence of the feature measured. Abbreviations: WGA, wheat germ agglutinin; * p < 0.05 compared to saline-challenged pigs. For panels E-F, data were assessed with a non-parametric Mann-Whitney test. All data are shown as mean ± SEM.

The presence of mucus sheets in the airways of pigs challenged with bethanechol suggested MUC5AC and MUC5B abundance might be increased. Because abnormal mucin protein concentrations have been implicated in chronic airway disease [33], we stained tracheal crosssections using antibody-specific labeling and performed signal intensity analyses [18, 34]. No significant differences were found in MUC5AC or MUC5B signal intensity within and on the tracheal surface between treatment groups (Figure 2A, 2B). Similarly, no differences in MUC5AC or MUC5B abundance was observed in the submucosal glands (Figure 2C, 2D). As a secondary approach, we also examined MUC5AC and MUC5B concentrations in the bronchoalveolar lavage fluid. Similar to the results obtained from antibody-labeling, we observed no differences between treatment groups (Figure 2E, 2F).

**Figure 2.**
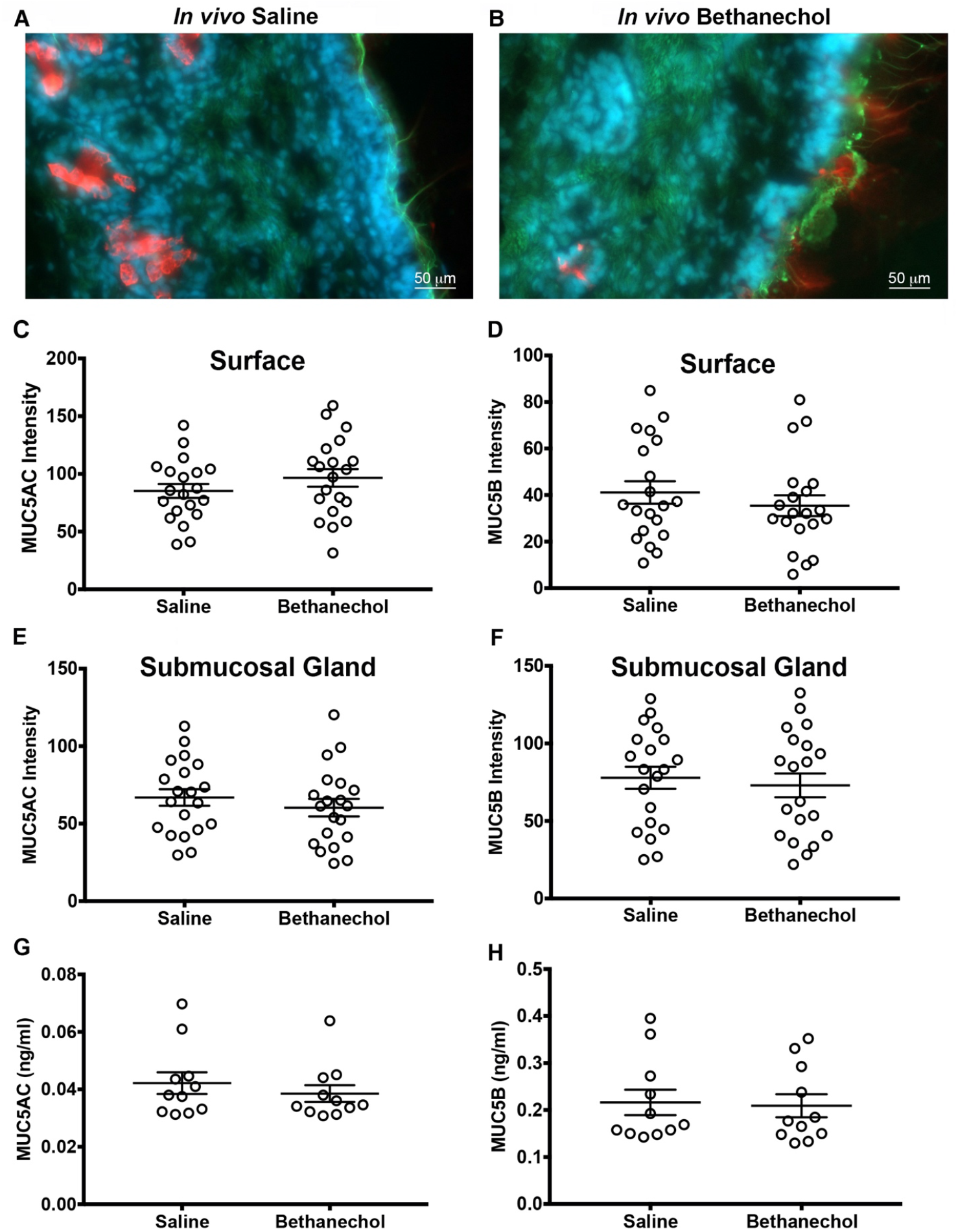
Tracheal mucin staining and bronchoalveolar concentrations in pigs challenged with bethanechol. Representative antibody-labeling of mucin 5AC (green) and mucin 5B (red) in tracheal cross sections from piglets challenged with intra-airway saline (**A**) or intra-airway bethanechol (**B**). Nuclei are shown in blue (Hoechst). MUC5AC and MUC5B staining signal intensity of the surface epithelia (**C, D**) and submucosal gland (**E, F**). Bronchoalveolar lavage fluid concentrations of MUC5AC (**G**) and MUC5B (**H**). For panels C-F, n = 20 saline-challenged piglets (10 females, 10 males), n = 20 bethanechol-challenged pigs (10 females, 10 males). For panels G and H, n = 11 saline-challenged pigs (5 females, 6 males) and n = 11 bethanechol-challenged pigs (5 females, 6 males). Abbreviations: MUC5B, mucin 5B; MUC5AC, mucin 5AC. All data are shown as mean ± SEM.

The lack of an effect of bethanechol treatment on MU5AC and MUC5B protein abundance suggested that enhanced concentrations of MUC5B and/or MUC5AC were not responsible for the formation of mucus sheets. Thus, we examined known modulators of mucus biophysical properties, namely fluid and ion transport, in primary cultures of porcine airway epithelia treated with bethanechol or saline control. We measured the apical surface fluid meniscus using previously published methods [19, 30]. Baseline volumes were consistent with previously published values (Figure 3A) [19, 30]. In saline-treated cultures, basolateral carbachol induced a significant increase in fluid secretion that was detected 30 minutes post-stimulation (Figure 3A, 3B). Carbachol was without effect in cultures treated with bethanechol (Figure 3A, 3B).

**Figure 3.**
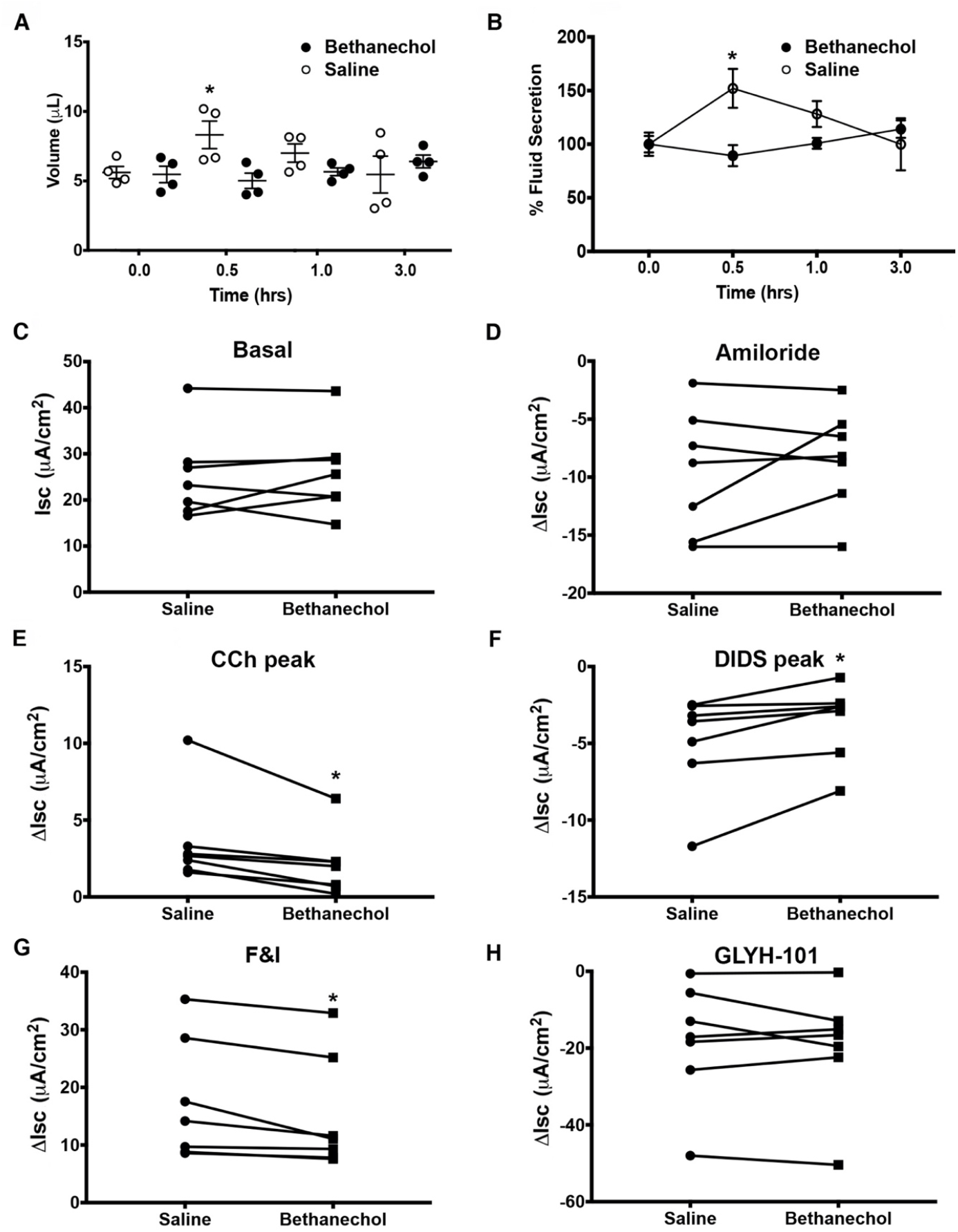
Fluid secretion and ion transport in cultured airway epithelia treated with bethanechol. (**A**) Apical fluid secretion in response to 100 μM basolateral carbachol in control and bethanechol-treated cultures. (**B**) Normalized fluid secretion as a percent of baseline. (C) Basal short circuit current (Isc) measurements in primary airway epithelia. Averaged delta (Δ) Isc measurements shown for: (**D**) 100 μM apical amiloride (AMIL); (**E**) 100 μM basolateral carbachol (CCh); (**F**) 100 μM apical DIDS inhibition from peak; (**G**) apical 10 μM forskolin and 100 μM IBMX (F&I); (**H**) 100 μM of apical GLYH-101. For panels A and B, n = 4 cultures (2 female and 2 male) per saline-treated and bethanechol treated groups. For panels C-H, n = 7 cultures (3 female and 4 males) per saline-treated and bethanechol-treated groups. Abbreviations: hour (hr). Data in panels A and B were assessed using a repeated measures ANOVA followed by a Sidak post hoc test. Data in panels C-H were assessed with paired Student’s t-test. * = p < 0.05 compared saline-challenged control. All data are shown as mean ± SEM.

Since ion transport regulates fluid secretion, we also measured short circuit current (Isc) in bethanechol and saline-treated cultures. Average basal Isc were not different between treatment groups (Figure 3C). Similarly, amiloride-sensitive ion transport was also not different (Figure 3D). However, both cholinergic-mediated Cl^-^ secretion (Figure 3E, 3F) and forskolin and IBMX (F&I)-mediated Cl^-^ secretion (Figure 3G) were decreased. Application of apical GlyH-101, a CFTR inhibitor [35], decreased F&I-mediated anion secretion in both treatment groups (Figure 3H). However, the extent of GlyH-101-inhibited current was not different between treatment groups (Figure 3H).

Impaired mucociliary transport is often a consequence of abnormal mucus secretion properties. Thus, we examined mucociliary transport *ex vivo* and using live imaging and computer-assisted particle tracking [18] (Figure 4). No differences were observed in mean or maximum speed of mucus transport (Figure 4C, 4D). Because speed is equal to distance over time, we also examined computer assigned particle track length and found it was not different (Figure 4E). Lastly, at the conclusion of the experiment, we also measured the signal intensity of fluorescently labeled mucus on the airway surface (Figure 4F). No differences between treatment groups were observed.

**Figure 4.**
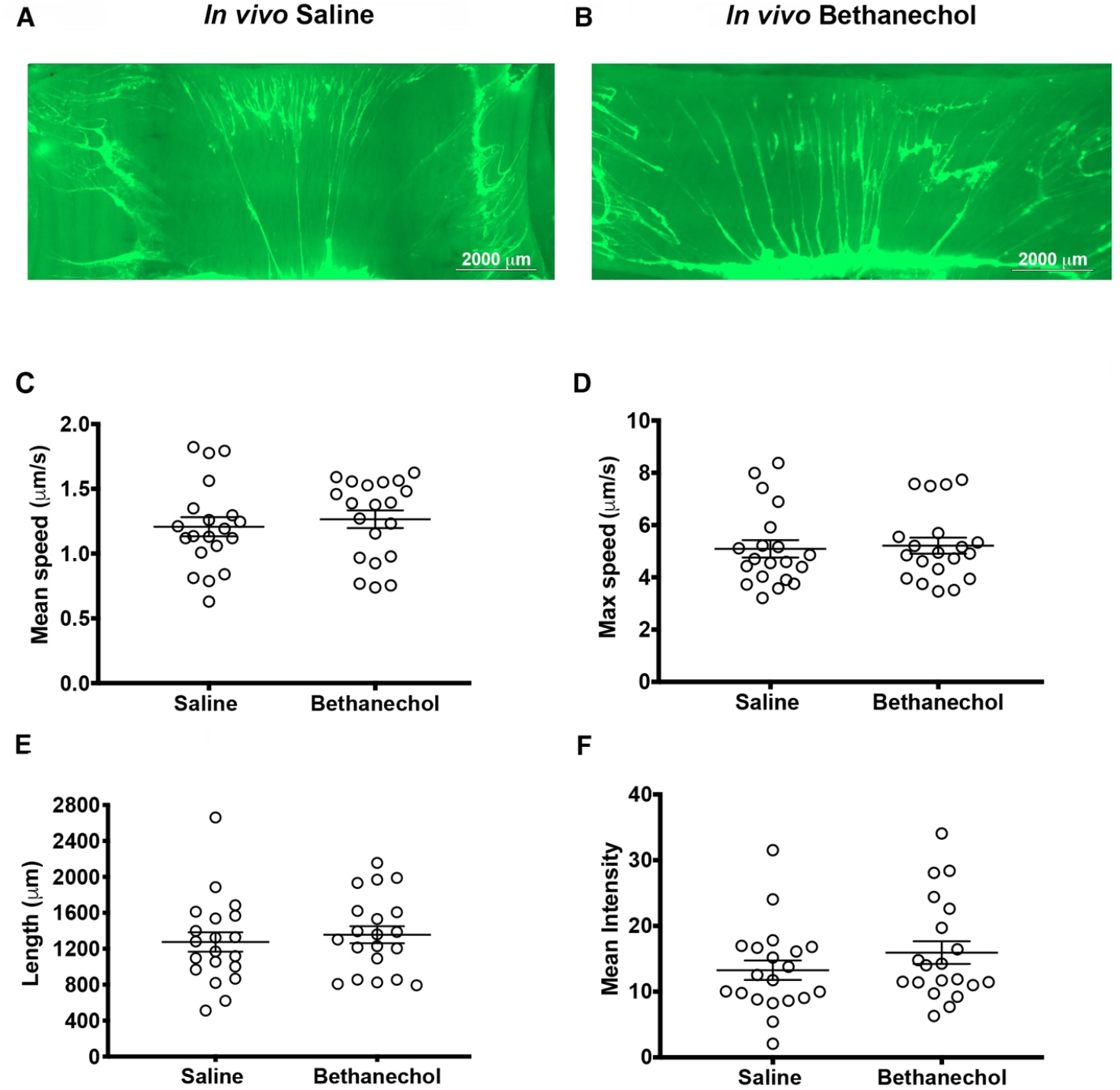
Mucus transport in piglet airways. Representative images of an *ex vivo* piglet tracheas stimulated with methacholine. Mucus is visualized in real-time with fluorescent nanospheres (bright green). Mucus often forms strands. Images of tracheas are at the conclusion of the experiment post methacholine stimulation in saline-challenged (**A**), and bethanechol-challenged (**B**) piglet airways. (**C**) Mean mucus transport speed. (**D**) Max mucus transport speed. (**E**) Computer assigned particle-track length. (**F**) Quantification of the signal intensity of fluorescently labeled mucus on the airway surface at the conclusion of the experiment. n = 20 saline-challenged piglets (10 females, 10 males), n = 20 bethanechol-challenged pigs (10 females, 10 males). Data were assessed with unpaired Student’s t-test. Mean ± S.E.M shown.

## Discussion

Mucus abnormalities characterize several airway diseases [36–38]. Although enhanced cholinergic transmission is thought to drive mucus abnormalities in inflamed airways, whether augmented cholinergic stimulation alone leads to lasting changes in mucus properties is largely unknown. Thus, we interrogated the sustained effect of cholinergic stimulation on mucus properties in neonatal piglets challenged with intra-airway bethanechol or saline control. Bethanechol elicited abnormal airway mucus characterized by persistent sheets, which were not present in saline treated controls. This phenotype was independent of differences in MUC5AC or MUC5B concentration measured in the bronchoalveolar lavage fluid or within the tracheal epithelia. Thus, these data suggest that cholinergic transmission *per se* is capable of inducing durable changes to mucus and fluid secretion.

Although bethanechol stimulation was associated with mucus sheet formation and diminished fluid secretion and Cl^-^ transport, no decrements in mucociliary transport were observed. This finding suggests that additional factors, such as robust inflammation and/or altered concentrations of mucins[39, 40], are required for mucociliary transport defects [41]. Another consideration is that submucosal gland fluid secretion and/or augmentation of ciliary beat frequency[42] compensate for the diminished fluid secretion mediated by surface epithelia, masking the detection of a defect. Conversely, it is interesting to speculate that cholinergic-mediated defects in fluid and mucus secretion might be compounded by innate deficits in fluid secretion in diseases such as CF [43, 44]. Indeed, mucociliary transport defects in CF pigs were prominent upon methacholine stimulation [24]. Lastly, since our studies were conducted in the absence of inflammation, it is difficult to predict how the current findings translate and map onto a background of inflammation.

Previous studies have demonstrated that anticholinergics reduce and delay exacerbations in COPD [45, 46]. Similar findings have been observed in asthma [15, 47]. Although our data support the importance of anticholinergics, they also raise a quandary. Specifically, anticholinergics are expected to decrease airway fluid secretion [48]. Yet we found that augmented cholinergic transmission also decreased fluid secretion. Perhaps this can be interpreted to indicate that the benefit of reducing mucus secretion with anticholinergics exceeds the potential consequence of decreasing fluid secretion. This interpretation also is consistent with our findings showing a lack of effect of decreased fluid secretion on mucociliary transport.

Our study has limitations. It is challenging to predict the exact temporal dynamics of cholinergic stimulation using aerosolized bethanechol. Variation in bethanechol distribution and clearance may contribute to heterogeneity in observed mucus stasis phenotypes. Moreover, we did not investigate mucus biophysical or rheological properties. We also did not measure airway surface liquid height *in vivo*, nor ciliary beat frequency. However, an advantage of bethanechol is its relatively long pharmacological activity due to the inability of hydrolyzation by cholinesterases. Also, given its solubility in water, bethanechol likely persists in the airway surface liquid periciliary layer, putatively allowing more sustained stimulation than systemic administration. Additionally, airway delivery likely mitigates undesired cholinergic stimulation effects on non-pulmonary organs and allowing investigation of airway surface properties.

In summary, augmented cholinergic transmission produces sustained mucus abnormalities characterized by persistent mucus films, decreased fluid secretion, and diminished Cl^-^ transport. Thus, it is possible that cholinergic transmission might perpetuate mucus abnormalities observed in several airway diseases during periods of exacerbations.

## Availability of data and material

No significant retained intellectual property associated with this manuscript.

## Acknowledgements

We thank Joshua Dadural, Shin-Ping Kuan, Kevin Vogt, and Eda Eken for excellent technical assistance. This study was in part funded by the National Institutes of Health: R00HL119560, OT2OD026582 (PI, LRR), OT2OD023854 (Co-I, LRR), and 5T32EB021955 (MJH).

## Author contributions statement

LRR and MJH conceived the study. LRR, MJH, and ENC wrote the manuscript. YSJL, ENC, MVG, KRA, VS, LB, MS and LRR performed experiments. LRR, ENC, LB, KRA, VS, YSJL and MVG performed data analysis. All authors reviewed and edited the manuscript.

## References

1. Dunican EM, Elicker BM, Gierada DS, Nagle SK, Schiebler ML, Newell JD, Raymond WW, Lachowicz-Scroggins ME, Di Maio S, Hoffman EA, et al: Mucus plugs in patients with asthma linked to eosinophilia and airflow obstruction. J Clin Invest 2018, 128:997–1009.

2. Burgel PR, Montani D, Danel C, Dusser DJ, Nadel JA: A morphometric study of mucins and small airway plugging in cystic fibrosis. Thorax 2007, 62:153–161.

3. Williams OW, Sharafkhaneh A, Kim V, Dickey BF, Evans CM: Airway mucus: From production to secretion. Am J Respir Cell Mol Biol 2006, 34:527–536.

4. Thornton DJ, Sheehan JK, Carlstedt I: Heterogeneity of mucus glycoproteins from cystic fibrotic sputum. Are there different families of mucins? Biochem J 1991, 276 (Pt 3):677–682.

5. Turkovic L, Caudri D, Rosenow T, Hall G, Stick S: Presence of mucus plugging is predictive of long term lung function in children with cystic fibrosis. European Respiratory Journal 2017, 50: OA4401.

6. Pearson MG, Ryland I, Harrison BD: Comparison of the process of care of acute severe asthma in adults admitted to hospital before and 1 yr after the publication of national guidelines. Respir Med 1996, 90:539–545.

7. Barr RG, Woodruff PG, Clark S, Camargo CA, Jr.: Sudden-onset asthma exacerbations: clinical features, response to therapy, and 2-week follow-up. Multicenter Airway Research Collaboration (MARC) investigators. Eur Respir J 2000, 15:266–273.

8. Fahy JV, Dickey BF: Airway mucus function and dysfunction. N Engl J Med 2010, 363:2233–2247.

9. Shaykhiev R: Emerging biology of persistent mucous cell hyperplasia in COPD. Thorax 2019, 74:4–6.

10. Canning BJ: Reflex regulation of airway smooth muscle tone. J Appl Physiol (1985) 2006, 101:971–985.

11. Wine JJ: Parasympathetic control of airway submucosal glands: central reflexes and the airway intrinsic nervous system. Auton Neurosci 2007, 133:35–54.

12. Jacoby DB, Xiao HQ, Lee NH, Chan-Li Y, Fryer AD: Virus- and interferon-induced loss of inhibitory M-2 muscarinic receptor function and gene expression in cultured airway parasympathetic neurons. Journal of Clinical Investigation 1998, 102:242–248.

13. Rios JD, Zoukhri D, Rawe IM, Hodges RR, Zieske JD, Dartt DA: Immunolocalization of muscarinic and VIP receptor subtypes and their role in stimulating goblet cell secretion. Invest Ophthalmol Vis Sci 1999, 40:1102–1111.

14. Scullion JE: The development of anticholinergics in the management of COPD. Int J Chron Obstruct Pulmon Dis 2007, 2:33–40.

15. Gosens R, Gross N: The mode of action of anticholinergics in asthma. Eur Respir J 2018, 52.

16. Reznikov LR, Meyerholz DK, Kuan SP, Guevara MV, Atanasova KR, Abou Alaiwa MH: Solitary Cholinergic Stimulation Induces Airway Hyperreactivity and Transcription of Distinct Pro-inflammatory Pathways. Lung 2018, 196:219–229.

17. Rodriguez E, Bullard CM, Armani MH, Miller TL, Shaffer TH: Comparison Study of Airway Reactivity Outcomes due to a Pharmacologic Challenge Test: Impulse Oscillometry versus Least Mean Squared Analysis Techniques. Pulm Med 2013, 2013:618576.

18. Liao YSJ, Kuan SP, Guevara MV, Collins EN, Atanasova KR, Dadural JS, Vogt KM, Schurmann V, Bravo L, Eken E, et al: Acid exposure disrupts mucus secretion and impairs mucociliary transport in neonatal piglet airways. Am J Physiol Lung Cell Mol Physiol 2020.

19. Kuan SP, Liao YJ, Davis KM, Messer JG, Zubcevic J, Aguirre JI, Reznikov LR: Attenuated Amiloride-Sensitive Current and Augmented Calcium-Activated Chloride Current in Marsh Rice Rat (Oryzomys palustris) Airways. iScience 2019, 19:737–748.

20. Reznikov LR, Liao YSJ, Gu T, Davis KM, Kuan SP, Atanasova KR, Dadural JS, Collins EN, Guevara MV, Vogt K: Sex-specific airway hyperreactivity and sex-specific transcriptome remodeling in neonatal piglets challenged with intra-airway acid. Am J Physiol Lung Cell Mol Physiol 2019, 316:L131–L143.

21. Reznikov LR, Meyerholz DK, Adam RJ, Abou Alaiwa M, Jaffer O, Michalski AS, Powers LS, Price MP, Stoltz DA, Welsh MJ: Acid-Sensing Ion Channel 1a Contributes to Airway Hyperreactivity in Mice. PLoS One 2016, 11:e0166089.

22. Ostedgaard LS, Moninger TO, McMenimen JD, Sawin NM, Parker CP, Thornell IM, Powers LS, Gansemer ND, Bouzek DC, Cook DP, et al: Gel-forming mucins form distinct morphologic structures in airways. Proc Natl Acad Sci U S A 2017, 114:6842–6847.

23. Abdullah LH, Wolber C, Kesimer M, Sheehan JK, Davis CW: Studying mucin secretion from human bronchial epithelial cell primary cultures. Methods Mol Biol 2012, 842:259–277.

24. Hoegger MJ, Fischer AJ, McMenimen JD, Ostedgaard LS, Tucker AJ, Awadalla MA, Moninger TO, Michalski AS, Hoffman EA, Zabner J, et al: Impaired mucus detachment disrupts mucociliary transport in a piglet model of cystic fibrosis. Science 2014, 345:818–822.

25. Jaqaman K, Loerke D, Mettlen M, Kuwata H, Grinstein S, Schmid SL, Danuser G: Robust single-particle tracking in live-cell time-lapse sequences. Nat Methods 2008, 5:695–702.

26. Karp PH, Moninger TO, Weber SP, Nesselhauf TS, Launspach JL, Zabner J, Welsh MJ: An in vitro model of differentiated human airway epithelia. Methods for establishing primary cultures. Methods Mol Biol 2002, 188:115–137.

27. Chen JH, Stoltz DA, Karp PH, Ernst SE, Pezzulo AA, Moninger TO, Rector MV, Reznikov LR, Launspach JL, Chaloner K, et al: Loss of anion transport without increased sodium absorption characterizes newborn porcine cystic fibrosis airway epithelia. Cell 2010, 143:911–923.

28. Stoltz DA, Rokhlina T, Ernst SE, Pezzulo AA, Ostedgaard LS, Karp PH, Samuel MS, Reznikov LR, Rector MV, Gansemer ND, et al: Intestinal CFTR expression alleviates meconium ileus in cystic fibrosis pigs. J Clin Invest 2013, 123:2685–2693.

29. He Q, Halm ST, Zhang J, Halm DR: Activation of the basolateral membrane Cl-conductance essential for electrogenic K+ secretion suppresses electrogenic Cl-secretion. Exp Physiol 2011, 96:305–316.

30. Harvey PR, Tarran R, Garoff S, Myerburg MM: Measurement of the airway surface liquid volume with simple light refraction microscopy. Am J Respir Cell Mol Biol 2011, 45:592–599.

31. Wang R, Ware JH: Detecting moderator effects using subgroup analyses. Prev Sci 2013, 14:111–120.

32. Ermund A, Meiss LN, Dolan B, Bahr A, Klymiuk N, Hansson GC: The mucus bundles responsible for airway cleaning are retained in cystic fibrosis and by cholinergic stimulation. Eur Respir J 2018, 52.

33. Lachowicz-Scroggins ME, Yuan S, Kerr SC, Dunican EM, Yu M, Carrington SD, Fahy JV: Abnormalities in MUC5AC and MUC5B Protein in Airway Mucus in Asthma. Am J Respir Crit Care Med 2016, 194:1296–1299.

34. Esther CR, Jr., Muhlebach MS, Ehre C, Hill DB, Wolfgang MC, Kesimer M, Ramsey KA, Markovetz MR, Garbarine IC, Forest MG, et al: Mucus accumulation in the lungs precedes structural changes and infection in children with cystic fibrosis. Sci Transl Med 2019, 11.

35. Muanprasat C, Sonawane ND, Salinas D, Taddei A, Galietta LJ, Verkman AS: Discovery of glycine hydrazide pore-occluding CFTR inhibitors: mechanism, structure-activity analysis, and in vivo efficacy. J Gen Physiol 2004, 124:125–137.

36. Pezzulo AA, Tang XX, Hoegger MJ, Alaiwa MH, Ramachandran S, Moninger TO, Karp PH, Wohlford-Lenane CL, Haagsman HP, van Eijk M, et al: Reduced airway surface pH impairs bacterial killing in the porcine cystic fibrosis lung. Nature 2012, 487:109–113.

37. Birket SE, Davis JM, Fernandez CM, Tuggle KL, Oden AM, Chu KK, Tearney GJ, Fanucchi MV, Sorscher EJ, Rowe SM: Development of an airway mucus defect in the cystic fibrosis rat. JCI Insight 2018, 3.

38. Abou Alaiwa MH, Beer AM, Pezzulo AA, Launspach JL, Horan RA, Stoltz DA, Starner TD, Welsh MJ, Zabner J: Neonates with cystic fibrosis have a reduced nasal liquid pH; a small pilot study. J Cyst Fibros 2014, 13:373–377.

39. Evans CM, Kim K, Tuvim MJ, Dickey BF: Mucus hypersecretion in asthma: causes and effects. Curr Opin Pulm Med 2009, 15:4–11.

40. Roy MG, Livraghi-Butrico A, Fletcher AA, McElwee MM, Evans SE, Boerner RM, Alexander SN, Bellinghausen LK, Song AS, Petrova YM, et al: Muc5b is required for airway defence. Nature 2014, 505:412–416.

41. Atanasova KR, Reznikov LR: Strategies for measuring airway mucus and mucins. Respir Res 2019, 20:261.

42. Zagoory O, Braiman A, Priel Z: The mechanism of ciliary stimulation by acetylcholine: roles of calcium, PKA, and PKG. J Gen Physiol 2002, 119:329–339.

43. Joo NS, Cho HJ, Khansaheb M, Wine JJ: Hyposecretion of fluid from tracheal submucosal glands of CFTR-deficient pigs. J Clin Invest 2010, 120:3161–3166.

44. Joo NS, Irokawa T, Robbins RC, Wine JJ: Hyposecretion, not hyperabsorption, is the basic defect of cystic fibrosis airway glands. J Biol Chem 2006, 281:7392–7398.

45. Casaburi R, Mahler DA, Jones PW, Wanner A, San PG, ZuWallack RL, Menjoge SS, Serby CW, Witek T, Jr.: A long-term evaluation of once-daily inhaled tiotropium in chronic obstructive pulmonary disease. Eur Respir J 2002, 19:217–224.

46. Vincken W, van Noord JA, Greefhorst AP, Bantje TA, Kesten S, Korducki L, Cornelissen PJ, Dutch/Belgian Tiotropium Study G: Improved health outcomes in patients with COPD during 1 yr’s treatment with tiotropium. Eur Respir J 2002, 19:209–216.

47. Quirce S, Dominguez-Ortega J, Barranco P: Anticholinergics for treatment of asthma. J Investig Allergol Clin Immunol 2015, 25:84–93; quiz 94-85.

48. Khansaheb M, Choi JY, Joo NS, Yang YM, Krouse M, Wine JJ: Properties of substance P-stimulated mucus secretion from porcine tracheal submucosal glands. Am J Physiol Lung Cell Mol Physiol 2011, 300:L370–379.

